# Distinct projection patterns of different classes of layer 2 principal neurons in the olfactory cortex

**DOI:** 10.1101/113134

**Authors:** Camille Mazo, Julien Grimaud, Venkatesh N. Murthy, C. Geoffrey Lau

**Affiliations:** Department of Molecular and Cellular Biology and Center for Brain Science, Harvard University, Cambridge, MA, USA; ENS de Cachan, Université Paris-Saclay, F-94235, Cachan, France; Department of Biomedical Sciences and Centre for Biosystems, Neuroscience, and Nanotechnology, City University of Hong Kong, 83 Tat Chee Avenue, Kowloon, Hong Kong

## Abstract

Processing of olfactory information in the anterior piriform cortex (APC) is widely thought to be non-topographic due to distributed projections from the olfactory bulb (OB). Layer 2 principal neurons in the APC can be divided into 2 subtypes: semilunar (SL) and superficial pyramidal (SP) cells. Although it is known that SL and SP cells receive differential inputs from the OB, little is known about their projection pattern to downstream structures, such as the posterior piriform cortex (PPC). Here we examined feedforward and feedback axonal projections of SL and SP cells using a combination of mouse genetics, and retrograde labeling. Retrograde tracing from the OB or PPC showed that the APC projects to these higher and lower brain regions mainly through layer 2b cells, while dual-labeling revealed a sizeable fraction of cells extending collaterals to both target regions. Furthermore, a transgenic mouse line specifically labeling SL cells showed that they send profuse axonal projections to olfactory cortical areas, but not to the OB. These findings support a model in which information flow from SL to SP cells and back to the OB forms a hierarchical feedback circuit whereas the two cell types process recurrent and feedforward information in a parallel manner.

## INTRODUCTION

Sensory perception emerges from the confluence of bottom-up and top-down inputs. In olfaction, feedback projections innervate the first brain relay for information processing: the olfactory bulb (OB). The OB receives information from olfactory receptor neurons, each bearing a single odorant receptor but expressing ~1,000 odorant receptors altogether in mice ^1,2^. Every sensory neuron expressing the same receptor converges to two glomeruli within each OB ^3^, where they directly synapse onto apical dendrites of OB principal cells, mitral and tufted cells, thereby forming an odotopic map. A given OB principal cell sends its apical dendrite to a single glomerulus, while the populations of mitral and tufted cell multiplex odor information to a variety of higher brain regions, such as the anterior piriform cortex (APC) ^4–6^. The APC is the largest region of primary olfactory cortex. It is involved in odor identity encoding, and is thought to serve as a storage unit ^7,8^. Single piriform neurons receive convergent inputs from multiple glomeruli. At the population level, odor information in the APC is sparse, distributed, and lacks evident topographic organization ^9–15^. Odor information encoded by assemblies of APC cells is then transmitted to a variety of olfactory regions such as the anterior olfactory nucleus (AON), the posterior piriform cortex (PPC), the cortical amygdala (CoA), or the lateral entorhinal cortex (LEnt). This information is also sent to non-primarily olfactory brain regions such as the orbitofrontal cortex (OFC). However, little remains known about the organization of APC projection channels.

The APC is a paleocortex, therefore composed of three layers. From superficial to deep: layer 1 is the input layer, layer 2 contains densely packed principal cells, and layer 3 comprises a combination of principal cells and GABAergic neurons. Deep to layer 3 is the endopiriform cortex (EndoP), mainly populated with multipolar neurons. Furthermore, layer 2 can be divided into two sublayers, 2a being roughly the superficial half of layer 2, and 2b the deeper half. Afferent inputs from the OB form synapses mainly with the distal dendrites of layer 2 principal cells. However, the strength and connectivity of these first synapses appear to be cell-type specific: the semilunar (SL) cells in L2a receive stronger inputs while the superficial pyramidal (SP) cells in L2b receive weaker sensory inputs ^16,17^. Although recurrent network activity and connectivity in the APC is well-described ^8,16,18,19^, recent work by Choy et al (2015) has demonstrated that connectivity scheme is cell-type specific. SL cells make synapses onto layer 2b SP cells without forming recurrent synapses on to themselves, while SP cells are recurrently connected ^20^. Therefore, layer 2 is populated with a mix of principal cells, namely SL and SP cells ^17^, playing different roles in the synaptic processing of olfactory information ^16^.

Input processing and recurrent connectivity is well described in the APC. However, it is unclear which neuron types contribute to the numerous projections out of the APC. Reconstruction studies of individual neurons suggest that APC principal cells project axons to the OB, the AON, and to downstream olfactory regions such as the PPC, LEnt, and CoA ^21,22^. However, it is unclear how prevalent are these cells that project both anterogradely and retrogradely. Recent work ^23,24^ confirmed original findings from Haberly and Price (1978)^25^, showing that feedback fibers from the APC to the OB do not originate homogeneously from all layers but appear to come from layers 2b and layer 3. In the present work, we used tracing techniques with pure retrograde activity, viral labeling, as well as mouse genetics to dissect the contribution of APC to upstream or downstream projections, with an emphasis on layer 2 principal neuron populations. We found that layer 2b is the main source of both feedback and feedforward projections, and that a sizeable fraction of neurons send collaterals to both regions. In addition, we found that genetically labeled SL cells projects widely to olfactory areas, but not back to the olfactory bulb.

## RESULTS

### The distribution of APC cells projecting back to the OB is biased toward layer 2b

To analyze the anatomical distribution of APC neuronal somas projecting back to the OB, we injected retrograde tracers in the OB of mice and examined the location of back-labeled somas in a series of sagittal slices of the APC (Figure 1A). We targeted our injections to the granule cell layer of the OB because it has been shown to be the largest recipient of feedback fibers originating from the APC ^26–29^.

**Figure 1.**
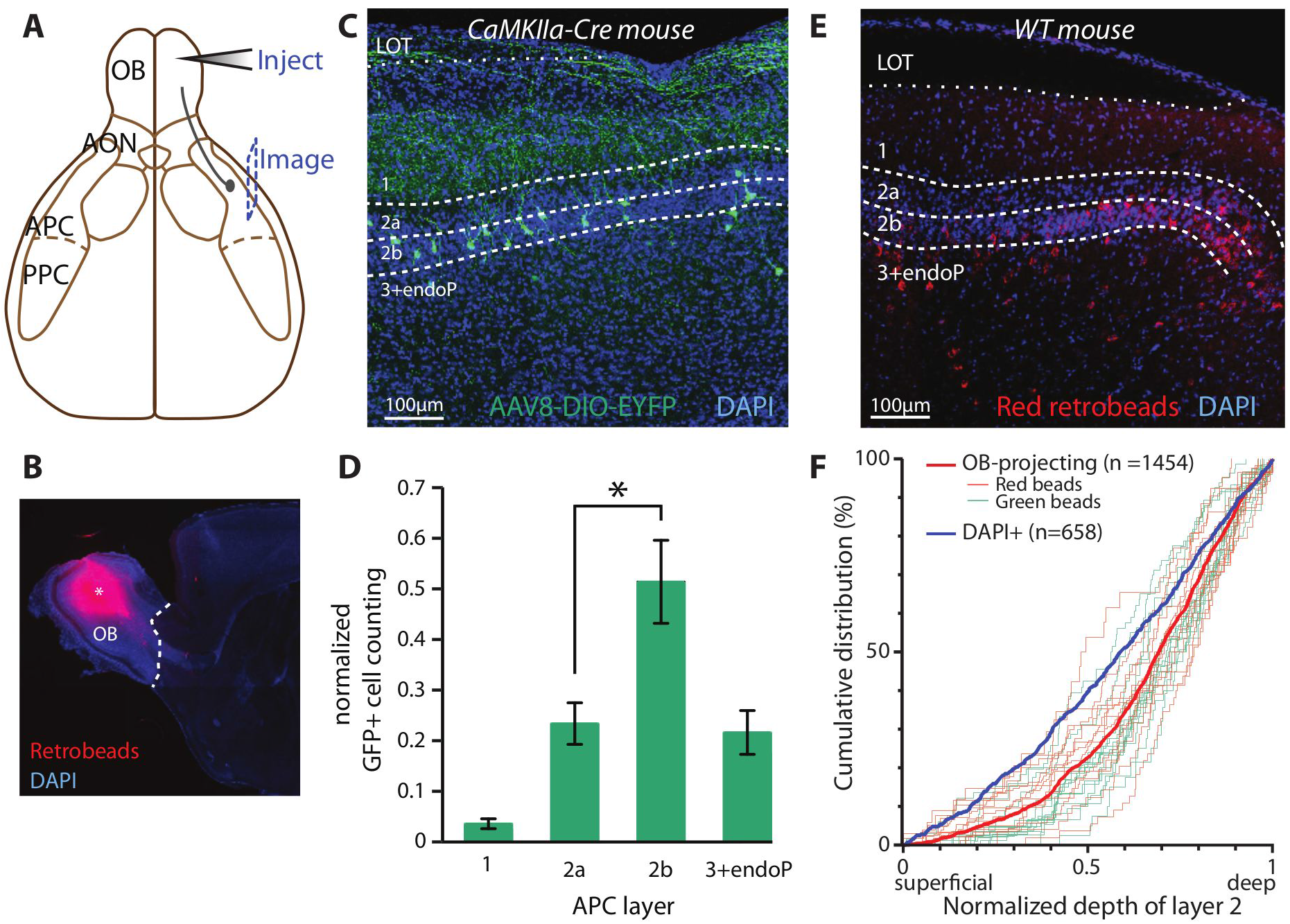
Layer 2b is the main source of feedback from the APC to the OB. A. Schematic representation of the injection site of the tracer (OB), and image obtained from sagittal APC slices. B. Injections were targeted to the granule cell layer. *, injection site. Red: tracer. Blue: DAPI. Similar results were obtained for the injections of AAV2/8-Flex-eYFP in CaMKIIa-Cre mice. C. Detection of neurons expressing eYFP in the APC of a CaMKIIa-Cre mouse, following OB injection of AAV2/8-Flex-eYFP. Green: eYFP. Blue: DAPI. D. Labeled neurons were more densely found in layer 2b than in layer 2a (Kruskal-Wallis test, p=0.022; post-hoc Mann-Whitney tests with Hochberg correction were performed to compare the layers two by two. All the tests were significant, except between layers 2a and 3+EndoP). E. Dense retrograde labeling of APC neurons was obtained after bead injection in the OB. Red: tracer. Blue: DAPI. F. Cumulative distribution of the OB-projecting cells within layer 2. On the x-axis, 0 indicates the border between layers 1 and 2a, while 1 is the limit between layers 2b and 3. The light green and red curves show the results from the counting obtained in a single optical section with green and red beads, respectively. The thick red trace plots the distribution of all the beads-labeled cells. The thick blue trace shows the biased distribution of the DAPI+ cells in layer 2. LOT: lateral olfactory tract.

Adeno-associated viruses (AAVs) have recently been shown to possess retrograde-labeling activity, especially when used with transgenic mice expressing Cre recombinase in specific neural populations ^30^. Thus, we injected Cre-dependent AAVs into OBs of CaMKIIa-Cre mice (Figure 1B). In our hands AAV capsid serotype 8 (AAV2/8-CAG-Flex-eGFP) worked the best to retrogradely label APC somas using this mouse line (Figure 1C; **Figure S1**). Following OB injections, retrogradely-labeled neurons were not evenly distributed across the different layers, as previously observed ^23–25^. 74.8 ± 6.1 % of the neurons were found in the layer 2, while 3.6 ± 1.0 % were located in layer 1, and 21.6 ± 4.3 % in layer 3 and the EndoP (Figure 1D), confirming the results from a similar study ^23^. Layer 2 can be further divided into 2 sublayers, namely layer 2a and 2b, from superficial to deep. When analyzing these two sublayers differentially, we observe that the distribution of OB-projecting neurons is biased toward layer 2b (23.4 ± 4.1 % for layer 2a versus 51.4 ± 0.8 % for layer 2b; Figure 1D), as reported by Diodato and colleagues (2016).

Viral infection could have influenced our observations, because the cell tropism of the virus may be biased toward specific neuron types. Therefore, in a new series of experiments, we injected either red or green retrobeads in the OB and analyzed the distribution of projecting neurons within layer 2 (Figure 1E). Here, we reported the continuous distribution of the retrogradely-labeled neurons as a function of the depth of layer 2 (0 being the superficial limit and 1 being the deep limit) and we estimated the depth profile of cell density within layer 2 using DAPI staining. Both red and green labeling led to a similar (p = 0.47, Kolmogorov-Smirnov test, n = 598 red bead- and n = 856 green bead-labeled cells), shifted distribution to the deepest part of layer 2, and therefore data were pooled (Figure 1F). The distribution of labeled cells in layer 2, compared to the cell distribution as measured using DAPI, was still biased toward the deepest part of layer 2 (p < 0.0001, Kolmogorov-Smirnov test, n = 1454 OB-projecting cells and n = 658 DAPI^+^ cells; Figure 1F), showing that the denser composition of layer 2b in cell bodies (comprising 59 ± 2% of L2 DAPI^+^ cells, *n* = 1372) is not sufficient to explain the shifted distribution of OB-projecting neurons within layer 2 of the APC.

It is noteworthy that bead injections in the OB led to labeling profiles comparable to what has previously been described in the literature, notably with ipsilateral labeling of somas in the AON pars principalis, but not in the AON pars externa; and contralateral labeling in AON pars principalis and AON pars externa, but also labeling of the horizontal limb of the diagonal band of Broca ^28,28,29,31–33^ (**Figure S2**).

Using both retrobeads and viral injections to label OB-projecting cells of the APC in a retrograde manner, we showed that the main OB-projecting population is found the layer 2b of the ipsilateral APC.

### The distribution of APC neurons projecting feedforward axons to the PPC is biased toward layer 2b

We then studied the distribution of the APC neurons targeting the PPC. Similarly to OB injections, retrobeads were injected in the PPC of mice and labeled somas were quantified in series of APC sagittal slices (Figure 2A-C). In contrast to the distribution of OB-projecting neurons (Figure 1D), the distribution of PPC-projecting neurons within the APC was bimodal, with a peak in layer 2 (peak at 74 % relative to depth of L2; 57% of cells) and a smaller peak in the EndoP (peak at 289 % of L2; 26% of cells); significantly fewer cells were to be found in layer 3 (17% of cells) (n = 225 cells, Figure 2C, D). These results corroborate the observations in an early work using horseradish peroxidase staining ^25^. Within layer 2, the distribution of the PPC-projecting population was skewed toward the deeper part of layer 2, and the ratio of projecting cells over the total number of DAPI+ cells was still biased toward layer 2b (p < 0.0001, Kolmogorov-Smirnov test, n = 499 PPC-projecting neurons, n = 658 DAPI+ cells; Figure 2E). Taken together, these data show that EndoP contains PPC-, but not OB-, projecting neurons, while layer 3 mostly contains OB-, but not PPC-, projecting neurons. In addition, the distribution of OB- and PPC-projecting neurons in layer 2 is skewed toward the deepest part of layer 2, layer 2b.

**Figure 2.**
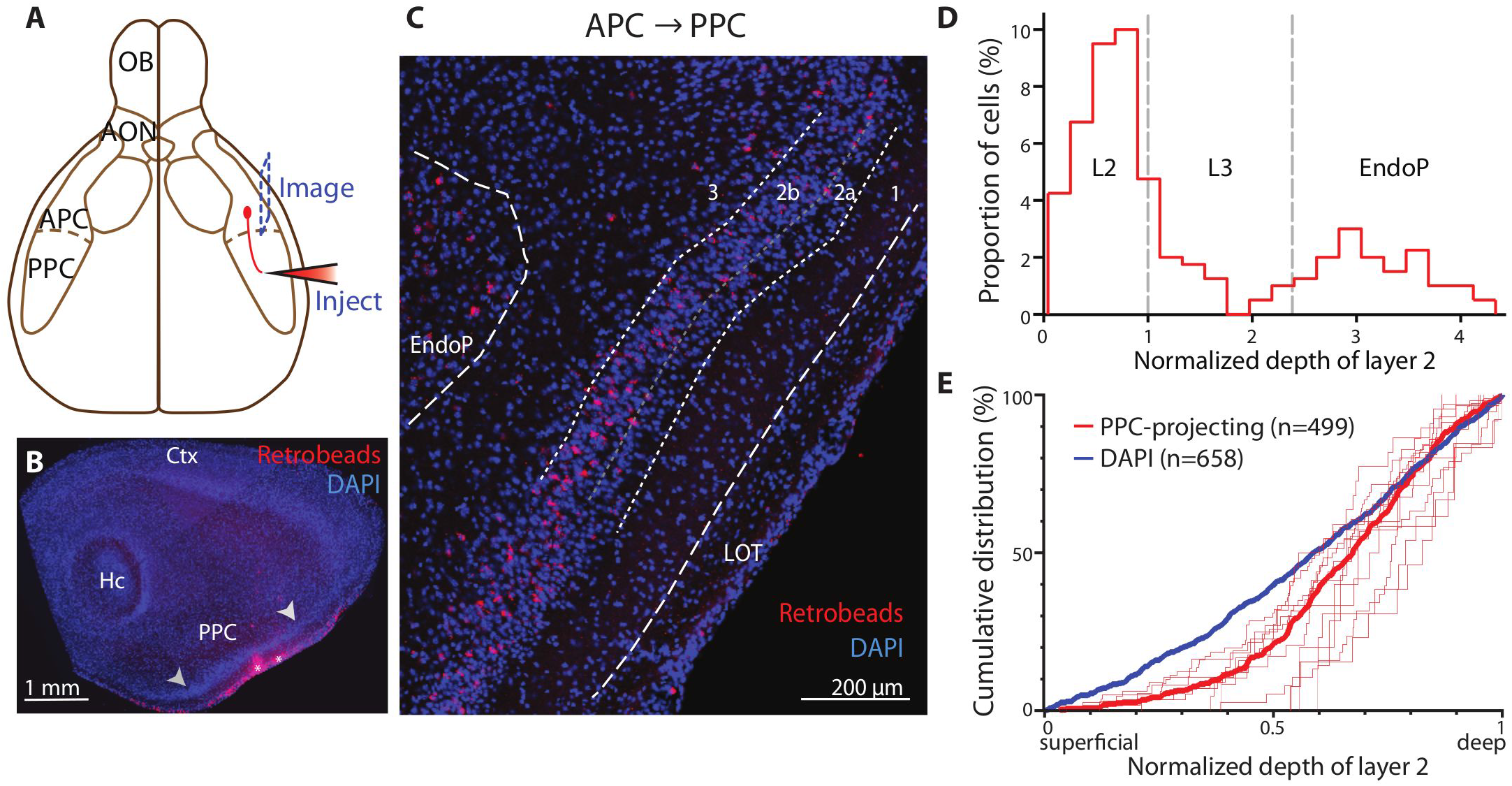
Layer 2b is also the predominant source of feedforward inputs from the APC to the PPC. A. Schematic representation of the injection site of the tracer (PPC), and image obtained from sagittal APC slices. B. Injections of tracer were targeted to the PPC. *, injection site. Arrowheads, PPC limits. Hc, hippocampus; Ctx, neocortex. *Inset,* zoom in the injection site. Scale bar is 200μm. C. Retrogradely labeled cells were found in APC sagittal slices. For B and C, red: tracer. Blue: DAPI. D. PPC-projecting cells were mainly located in deep layer 2 and in the EndoP. E. Within layer 2, projecting neuron distribution was skewed toward layer 2b. Thin red traces represent the counting for a single slice, while the thick red trace shows the distribution of all the counted cells. The thick blue trace is the distribution of the DAPI+ cells.

### OB- and PPC-projecting neurons are overlapping populations in layer 2 of the APC

We demonstrated that the OB- and PPC-projecting populations of the APC share dissimilar distribution patterns across the three layers and EndoP, but apparently similar patterns inside layer 2. We next asked whether the neurons from layer 2 projecting back to the OB and forward to the PPC belong to segregated or the same population of APC projecting cells, such that a single cell projects to both areas. Recent tracing work from Chen et al. (2014) ^34^ showed that APC axons projecting to two distinct areas of the OFC – namely the agranular insula and the lateral OFC – were spatially segregated within the APC, suggesting a topographical organization of the APC based on its output channels. To answer this question, injections of green retrobeads in the OB and red retrobeads in the PPC were performed in the same mice (Figure 3A). First, OB- and PPC-retrogradely labeled neurons were found to have indistinguishable distribution patterns in layer 2 (p > 0.05, Kolmogorov-Smirnov test, n = 782 and n = 499 OB- and PPC-projecting neurons, respectively; Figure 3B). Then, of the 783 retrogradely labeled neurons from the OB (green beads^+^), 91 of them (11.6%) were also projecting to the PPC (green and red beads^+^). Similarly, 91 of the 499 cells projecting to the PPC were found to project to the OB as well (18.2%; Figure 3C-E). The fact that we found non-zero percentages of dual-labeled cells shows that at least some of them project to both the PPC and OB.

**Figure 3.**
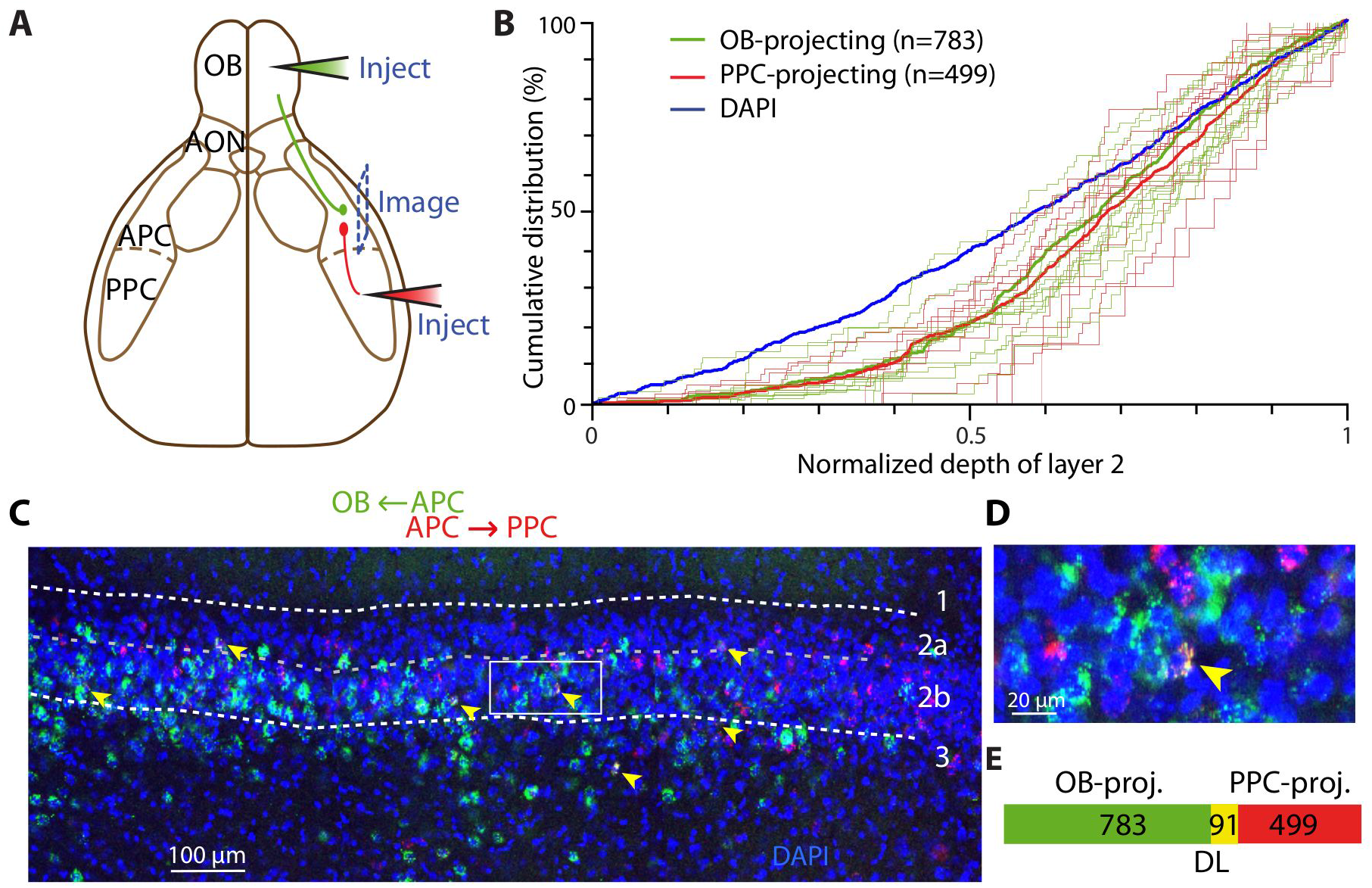
OB- and PPC-projecting cells are at least partially overlapping populations. A. Schematic representation of the dual injection strategy. Green beads were injected in the OB while red beads were injected in the PPC. Images were taken from the APC. B. OB- and PPC-projecting neurons share similar distribution within APC layer 2. The thin green and red traces represent quantifications from single optical sections. The thick green, red and blue traces are the distribution of all the OB-, PPC-projecting neurons and DAPI cells, respectively. Green: OB-projecting; Red: PPC-projecting. C. Example APC section with OB- (green) and PPC-projecting (red) neuron population, spatially overlapping. Arrowhead: dual-labeled neurons. D. Blow-up of the box in C., showing a dual-labeled neuron in layer 2b. E. Venn diagram representing the counted number of dual-labeled cells (DL) in each population.

However, several parameters may lead to an underestimation of the dual-labeled population. First, within the injection site, retrobeads labeled only a fraction of neurons that are actually projecting. To estimate the labeling efficiency, we sequentially injected identical amounts of green and red beads in the OB. Under these conditions, a large majority of labeled cells in the APC was double-labeled (90.5 − 92.7%, *n* = 205 − 210 cells; **Figure S3A**), similarly to a previous study ^35^. Thus, the co-labeling efficiency of green and red beads appears to be high and this factor only contributes weakly to the underestimation of dual-labeled population. Second, using our method, injections were spatially restricted to several hundred of micrometers (Figure 1A, 2A). Since we do not know how many APC neurons are actually projecting to the OB or PPC, we cannot estimate the fraction of actually projecting cells we are labeling using our beads injection protocol. Evidence for a certain degree of topography in OB-projection patterns ^24,36^ suggest that our restricted bead injection limits the number of possible retrogradely labeled neurons to those projecting to the precise injection locus. Notably, Matsutani (2010) ^36^ described patchy axon terminals in the OB. To measure whether restricted beads injection underestimated the fraction of projecting neurons, we injected green and red beads ~500μm apart within the OB and observed co-labeling of only a third of the projecting neurons (34.3 − 21.1%, *n* = 67 − 109 cells in 2 mice; **Figure S3B**),. Therefore, our results suggest that our injection protocol largely underestimates the number of neurons projecting to the OB or the PPC. As a result, the actual dual-labeled population is likely to be much larger than what is estimated here. We conclude from these dual-labeling experiments that a given layer 2 APC neuron is very likely to project to both the OB and the PPC.

### Outputs from SL cells within layer 2a are widely distributed in olfactory areas but do not project to the OB

We showed that the cell population of layer 2a in the APC is composed of at least a small fraction of neurons projecting to the OB or the PPC (Figure 1C-F, Figure 2C-E). Neurons in Layer 2a are mainly composed of SL cells, believed to be specialized in providing feedforward excitation to pyramidal cells of layer 2b and layer 3 ^16,17,20^. However, a single neuron originating in layer 2a can extend axons to multiple olfactory regions ^22^ and recent work combining genetics and tracing techniques showed that layer 2a cells project to distinct brain regions ^23^. Therefore, we took advantage of a genetic mouse line that specifically expresses the reporter protein mCitrine and tetracycline activator in a subpopulation of SL cells (48L mouse line, mCitrine expressed in 46 ± 2% of L2a Nissl-labeled cells)^20,37^ to investigate the projection pattern of SL cells in layer 2a (Figure 4A). Labeled mCitrine^+^ axons were found throughout upstream and downstream olfactory cortical areas including the AON, PPC, and CoA, while very few axons were labelled in the OB (Figure 4A), as previously reported for not genetically identified layer 2a neurons ^22,23^. A survey of various AAV serotypes revealed that AAV2/5 has the highest, while AAV2/8 has the lowest, labeling efficiency for SL cells (Figure S4). Notably, diluted AAV2/1 injections (AAV2/1-TRE::myr-mCherry; Figure 4B) in the APC led to sparse dual labeling of a SL cell subpopulation. Individual double-labeled axons projected as least 1 mm away from APC in both the anterior and posterior directions (Figure 4C). Furthermore, more concentrated injections of AAV2/5 in the same location labeled SL-mCitrine^+^ axons in various olfactory cortical regions including the AON, APC, and CoA (Figure 4C), but not in the OB. To directly visualize whether projections of SL cells spare the OB, we injected red retrobeads in the OB of 48L animals. Quantification reported a near absence of co-labeling between the genetic reporter of SL cells mCitrine and the injected retrobeads (3 dual-labeled cells out of 226 48L- and 225 bead-labeled cells, *n* = 3 slices, while PPC beads injection led to some dual-labeling; Figure 4D and **Figure S4B, C**). Taken together, our data show that SL cell population extends long-range projections to multiple olfactory cortical areas (such as the AON and the PPC), but sparing feedback to the OB.

**Figure 4.**
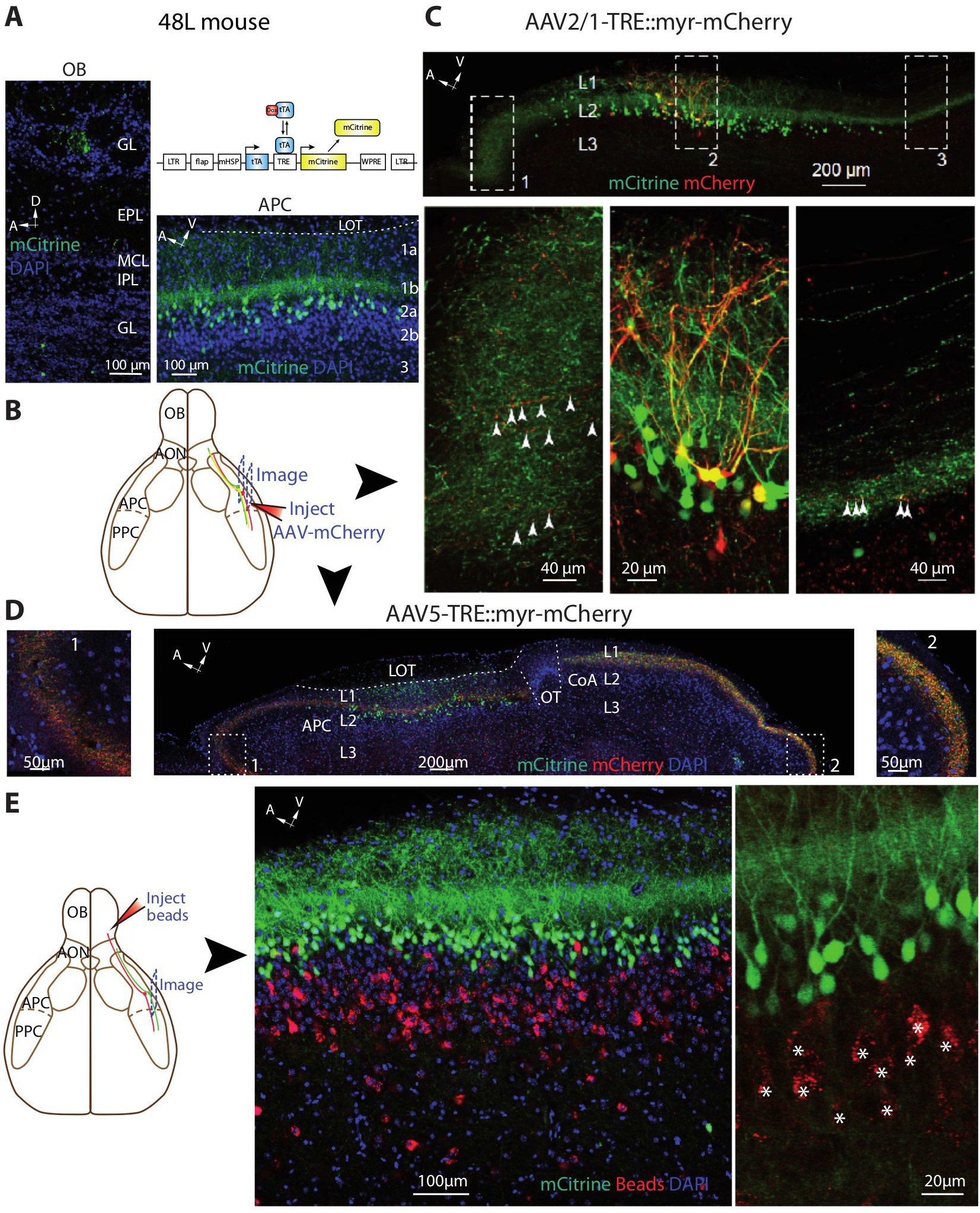
48L-labeled cells projects widely within the olfactory system, but not back to the OB. A. tTA binding to TRE drives mCitrine expression in a subset of SL cells (dox off system; *up right*). mCitrine^+^ cells were concentrated in layer 2a of the APC (*bottom right*). In the OB (*left*), some cells were observed in the glomerular and granule cells layers. Axons were rarely observed. GL: glomerular layer, EPL: external plexiform layer, MCL: mitral cell layer, IPL: internal plexiform layer, GCL: granule cell layer. B. AAVs expressing mCherry were injected in the APC to yield sparse dual-labeling and identification of dual-labeled axons away from the injection site. C. AAV2/1-TRE::myr-mCherry injection in the APC led to sparse dual-labeling of mCitrine^+^ cells (box 2). Dual-labeled axons were found several hundred μm away from the injection site in the sagittal plane, in the dorsal (box 1) and ventral APC (box 2). D. AAV2/5-TRE::myr-mCherry injection in the APC labeled axons several hundred μm away from the injection site, in a parallel plane. Dual-labeled axons were found in the dorsal APC and in the CoA. OT: Olfactory Tubercle. For B and C, red: mCherry. Green: mCitrine. Blue: DAPI. E. Red bead injections in the OB of 48L mouse failed to labeled mCitrine^+^ cells, indicating that the genetically tagged subset of SL cells is not projecting back to the OB. Middle and right panels are extracted from different experiments. Stars in right panels indicate beads^+^ OB-projecting cells. Red: beads. Green: mCitrine. Blue: DAPI.

## DISCUSSION

In this study, we used injections of retrograde tracers in the OB and PPC of wild-type and transgenic mice to examine the distribution as well as the projections of principal neurons from the APC. We characterized the laminar distribution of projecting populations and identified a significant number of neurons dually projecting to the OB and the PPC. In addition, we showed that genetically labeled SL cells project to numerous brain regions, but not back to the OB. Not only do these findings bring new knowledge about how the APC broadcasts olfactory information to the brain, they also reveal new insights into olfactory coding.

We found that OB-projecting neurons were mostly present in layers 2 and 3 of the APC, while PPC-projecting neurons were found mainly in layer 2 of the APC and in the EndoP. Within layer 2, both OB- and PPC-projecting populations were largely skewed towards layer 2b. Recent work from Diodato and colleagues (2016) found a similar distribution of OB-projecting neurons, and also reported a layer 2b-biased distribution of APC neurons projecting to the medial prefrontal cortex. Layer 2 of the APC is composed of superficial layer 2a, mainly populated with SL cells, and deep layer 2b, mainly composed of SP cells - although a continuum exists between the two cell populations ^16^. SL cells receive strong bottom-up inputs from the OB and form little or no recurrent connections with local excitatory neurons ^16,17,20^. In contrast, SP cells receive stronger inputs from recurrent axons and project outside the APC ^16,17,38^. Our data and previous work show that the main projection channel of the APC indeed originates from layer 2b, presumably from SP cells. However, in this study, we also genetically labeled a significant proportion of projecting layer 2a cells that were shown previously to be SL cells based on morphological and electrophysiological characterization ^20^. Genetic labeling revealed that SL cells do send axonal projections to multiple brain regions. Consistent with this observation, single-cell tracing studies observed SL cell axons outside the APC ^22^. Moreover, retrograde tracer injections in the CoA or LEnt by Diodato and coworkers (2016) identified layer 2a as the main APC output channels to these brain regions. Therefore, it appears that SL and SP cells of layer 2 constitute two parallel output channels of the APC. It is possible that SL cells send to other olfactory regions odor information that has been only subject to moderate local processing whereas SP cells send more processed signals owing to its recurrent connections.

Strikingly, while the genetic labeling of SL cells revealed projection to a variety of brain regions, it failed to reveal significant feedback projections to the OB (Figure 4A). We did observe very few axons in the OB, but these projections were very sparse and likely originates locally from labeled cells in the OB (juxtaglomerular cells in the glomerular layer or cells in the granule cell layer ^20^). Although 48L mouse line labels only a sub-population of SL cells (~46 ± 2% of L2a Nissl-labeled cells)^20^, it appears that OB-projecting cells in layer 2 of the APC does not originate from SL cells. Here, we propose a circuit model where SL and SP cell projections are organized differently depending on whether these are feedback or feedforward motifs. As SL cells receive stronger inputs from the OB with little or no recurrent connections, they are the first processing station in the APC. Information is fed forward from SL cells to higher olfactory regions as well as to SP cells within the APC. For feedback information, however, an addition processing is required: SL to SP and SP to SP connections will dictate the kind of information sent back to the OB. Since SL cells do not project back to the OB but instead rely on SP cells to relay feedback information, this forms a hierarchical processing circuit. On the other hand, feedforward processing occurs in a parallel fashion for SL and SP outputs (Figure 2 and 4).

Within the APC, OB- and PPC-projecting neurons were found mainly in layer 2. Dual-labeling experiments showed that a significant fraction of OB- or PPC-projecting neurons actually project to both areas. Therefore, it appears that information emerging from overlapping populations, and thus similar odor stimuli, are sent back to the OB and forward to the PPC. In contrast, using a similar dual-tracing technique, Chen et al. (2014) identified distinct OFC-projecting neuronal populations in the APC, although spatially overlapping. Genetic analysis of different projecting populations ^23^ further shows that spatial segregation of projection neurons is not required for forming target-specific cell populations. Additional connectivity studies, which might benefit from tissue clearing techniques, are necessary to gain insight on whether APC outputs are predominantly multiplexed or rather parallelized into distinct channels. We believe that a better understanding on odor coding in the brain requires more information about the output organization of the APC. It is likely that the formation of odor percepts involves wide recruitment of multiple brain areas and intricate feedback and feedforward circuits.

## METHODS

### Animals

C57B1/6J, CaMK2a-Cre ^39^ and 48L mice (labeling SL cells) were used in this study. All experiments were performed in accordance with the guidelines set by the National Institutes of Health and approved by the Institutional Animal Care and Use Committee at Harvard University.

### Retrograde labeling

In this study, non-viral tracers (green and red fluorophore-coated latex beads; Lumafluor), and viral tracers were used to retrogradely label APC projecting neurons.

For OB retrograde labeling (Figures 1 and S1), the viruses used were: AAV2/1-CAG::hChR2(H134R)-mCherry, AAV2/8-CAG::ChR2-GFP, AAV2/9-CAG::ChR2-Venus, AAV2/8-CAG::Flex-EYFP, and AAV2/9-Flex-ChR2-eYFP. All viruses were purchased from the Penn Vector Core. Cre-dependent viruses were used in CamK2a-Cre mice while and non-Cre dependent viruses were injected in C57BL/6J mice. For APC injections in 48L mice (Figures 4 and S4), we used AAV2-TRE::myr-mCherry with capside serotype 2/1, 2/5 or 2/8 (Shima and Nelson, Brandeis).

Briefly, adult male mice (1 to 4 months old) were deeply anesthetized with a ketamine/xylazine mixture (100 mg/kg and 10 mg/kg, respectively; i.p., Webster) and placed in a stereotaxic apparatus. A small craniotomy was performed above the injection site and labeling solution was injected into the OB, APC or PPC using a glass pipette (OB injection coordinates: AP 1.2 mm anterior to the sinus posterior to the OB; ML: 1.1 mm; and 1 mm deep from the brain surface; volume injected: 50 nL with beads, 300 nL with viral solutions, otherwise stated in the article. APC: ML 2.5−2.7 mm, AP 1.3-1.5 mm, 3.5-3.6 deep from bregma, 100nL of viral solution. PPC; AP: 0.3-0.7 mm anterior to lambda, ML: 3.4-4.2 mm; 4.4-5.1 mm deep from the skull surface, 50nL of beads).

### Histology and cell counting

1 to 4 days after retrobead injections or two weeks after viral injections, mice were perfused intracardially with 4% v/v paraformaldehyde and brains were post-fixed in the same fixative overnight. 100 μm-thick brain sections were cut with a vibratome (Leica VT1000 S), rinsed in PBS, counterstained with the nuclear dye 4,6-diamidino-2-phenylindole (DAPI) and mounted on slides. Z-stack confocal images were taken with a Zeiss 780 confocal microscope. The size of pinhole was adjusted to yield optical slice depth of approximately 10 μm. Two to three slices per animal were taken (sagittal sections, 200μm apart) and fluorescent cells were manually counted with the Fiji plugin “Cell Counter” by Kurt de Vos (University of Sheffield) on a maximum intensity projection of the Z-stack.

For the histogram quantifications, distinct regions were defined manually; corresponding to layer 1, layer 2a, layer 2b, layer 3 and EndoP. The number of labeled cells in each region was counted, as well as the number of DAPI+ cells. For each layer, the percentage of labeled neurons was normalized by the number of DAPI+ cells.

For cumulative plots, the depth of the cells in layer 2 was determined by measuring the distance of a cell to the layer 1 / layer 2 border and reported to the depth of layer 2 in that region. DAPI+ cells were quantified on the binarized stack in the middle of the Z-stack. The mean of DAPI+ cells across experiments was used for normalization in bar graphs.

### Statistics

All the results are given as mean ± standard deviation of the mean. In all graphs the mean is represented. All the statistical tests were performed using commercial analysis software (Graphpad Prism) or custom script in Matlab with a 5% significance level.

## AUTHOR CONTRIBUTIONS

V.N.M. and C.G.L. supervised the project. C.M., J.G. and C.G.L. performed experiments, analyzed data, and prepared figures. All authors wrote, reviewed, and edited the manuscript. V.N.M. provided research funding.

